# Neurofibromin 1 mediates sleep depth in *Drosophila*

**DOI:** 10.1101/2022.09.15.508161

**Authors:** Elizabeth B. Brown, Jiwei Zhang, Evan Lloyd, Elizabeth Lanzon, Valentina Botero, Seth Tomchik, Alex C. Keene

**Affiliations:** Department of Biology, Texas A&M University, College Station, TX, USA, 77840; Jupiter Life Science Initiative, Florida Atlantic University, Jupiter, FL 33431; Department of Neuroscience and Pharmacology, University of Iowa Carver College of Medicine, Iowa City, IA

## Abstract

Neural regulation of sleep and metabolic homeostasis are critical in many aspects of human health. Despite extensive epidemiological evidence linking sleep dysregulation with obesity, diabetes, and metabolic syndrome, little is known about the neural and molecular basis for the integration of sleep and metabolic function. The RAS GTPase-activating gene *Neurofibromin* (*Nf1*) has been implicated in the regulation of sleep and metabolic rate, raising the possibility that it serves to integrate these processes, but the effects on sleep consolidation and physiology remain poorly understood. A key hallmark of sleep depth in mammals and flies is a reduction in metabolic rate during sleep. Here, we use indirect calorimetry to define the role of *Nf1* on sleep-dependent changes in metabolic rate. Flies lacking *Nf1* fail to suppress metabolic rate during sleep, raising the possibility that loss of *Nf1* prevents flies from integrating sleep and metabolic state. Sleep of *Nf1* mutant flies is fragmented with a reduced arousal threshold in *Nf1* mutants, suggesting *Nf1* flies fail to enter deep sleep. The effects of *Nf1* on sleep can be localized to a subset of neurons expressing the GABA receptor *Rdl*. Selective knockdown of *Nf1* in *Rdl*-expressing neurons increases gut permeability and reactive oxygen species (ROS) in the gut, suggesting a critical role for deep sleep in gut homeostasis. Together, these findings suggest *Nf1* acts in GABA-sensitive neurons to modulate sleep depth in *Drosophila*.

## Introduction

The functional and neural basis of sleep is highly conserved from invertebrates through mammals (Joiner, 2016; Keene & Duboue, 2018). In many cases, powerful genetics in relatively simple model systems, including the fruit fly, *Drosophila melanogaster*, have allowed for the identification of novel genes and neural mechanisms that have informed our understanding of human sleep (Allada & Siegel, 2008; Sehgal & Mignot, 2011). However, most work in these models have studied total sleep duration. Therefore, a lack of understanding of the mechanisms underlying sleep quality and broader changes in physiology associated with sleep in non-mammalian models represents a significant gap in our knowledge. In mammals, slow-wave sleep is associated with reduced metabolic rate (Allison & Cicchetti, 1976; Berger et al., 1988; Sharma & Kavuru, 2010). Growing evidence suggests that many physiological changes associated with mammalian sleep are conserved in flies, including a reduction in whole body metabolic rate (Alphen et al., 2013; Faville et al., 2015; Yap et al., 2017). The diverse physiological changes associated with sleep, including changes in body temperature, reduced metabolic rate, and synaptic homeostasis, are thought to be critical for sleep’s rejuvenate properties (Krueger et al., 2016; Tononi & Cirelli, 2014; Zielinski et al., 2016).

Flies, like mammals, exhibit distinct electrophysiological patterns that correlate with wake and rest (Nitz et al., 2002; Raccuglia et al., 2019; Yap et al., 2017). We have identified sleep-associated reductions in metabolic rate in flies that are consistent with those that occur in mammals (Stahl et al., 2017, 2018). In addition, flies display all the behavioral hallmarks of sleep, including an extended period of behavioral quiescence, rebound following deprivation, increased arousal threshold, and species-specific posture (Hendricks et al., 2000; Shaw et al., 2000). Behavioral tracking systems and software are available for high-throughput detection and analysis of fly sleep using infrared monitoring or video tracking (Garbe et al., 2015; Gilestro, 2012). Sleep in *Drosophila* is typically defined as 5 minutes or more of behavioral quiescence, as this correlates with other behavioral and physiological characteristics that define sleep (Alphen et al., 2013; Stahl et al., 2017; Yap et al., 2017). For example, sleep bouts lasting ~10 minutes or longer are associated with increased arousal threshold and low-frequency oscillations in brain activity. These findings are supported by computational analysis modeling sleep pressure (Alphen et al., 2013; Wiggin et al., 2020; Yap et al., 2017). These analyses suggest the presence of light and deep sleep in flies; however, the genetic and neural basis for these different types of sleep is poorly understood.

The *Nf1* gene encodes a large protein that functions as a negative regulator of Ras signaling and mediates pleiotropic cellular and organismal function (Gutmann et al., 2017; Martin et al., 1990). *Nf1* mutations in humans cause a disorder called Neurofibromatosis Type 1, characterized by benign tumors of the nervous system (neurofibromas), as well as increased susceptibility to neurocognitive deficits (e.g., attention-deficit/hyperactivity disorder, autism spectrum disorder, visuospatial memory impairments;Gutmann et al., 2017b). In addition, mutation of *Nf1* is associated with dysregulated sleep and circadian rhythms (Licis et al., 2013; Williams et al., 2001). *Drosophila* deficient for *Nf1* recapitulate many of these phenotypes and are widely used as a model to investigate the role of *Nf1* in regulation of cellular and neural circuit function (Walker et al., 2012). Furthermore, *Drosophila Nf1* mutations lead to dysregulated circadian function and shortened sleep (King et al., 2016; Williams et al., 2001). Here, we examine the effects of *Nf1* on sleep-dependent changes in metabolic rate and measures of sleep depth.

We find that flies lacking *Nf1* fail to suppress metabolic rate during prolonged sleep bouts, revealing a disruption of sleep-dependent changes in metabolic rate. Furthermore, multiple behavioral measurements suggest sleep depth is disrupted in *Nf1* mutant flies, including the presence of sleep fragmentation and reduced arousal threshold. Genetic and pharmacological analysis suggest *Nf1* modulates GABA signaling to regulate sleep depth and sleep-dependent changes in metabolic rate. Therefore, these findings suggest that *Nf1* is a critical regulator of sleep-metabolism interactions, and the conserved molecular and phenotypic nature of *Nf1* mutants raises the possibility that these findings may be relevant to the complex pathologies in humans afflicted with *Nf1*.

## Results

To examine the effects of *Nf1* on sleep and activity, we compared sleep of control flies to *Nf1*^P1^ mutants that harbor a near-total deletion in the *Nf1* locus (The et al., 1997). Sleep was reduced during the day and night in *Nf1*^P1^ mutants compared to controls (Fig 1A,B). Sleep duration in *Nf1*^P1^ heterozygous flies did not differ from controls, indicating that the phenotype is recessive (Fig 1A,B). The average number of sleep bouts was increased in *Nf1*^P1^ flies, while the average bout length was reduced compared to control and heterozygote flies, suggesting that loss of *Nf1* results in sleep fragmentation (Fig 1C-D). In addition to the loss of sleep, the average velocity of activity during waking periods (waking activity) is elevated, suggesting that loss of *Nf1*^P1^ also results in hyperactivity (Fig S1A). These findings suggest *Nf1* promotes sleep duration, consolidation of sleep bouts, and modulates waking activity.

**Figure 1.**
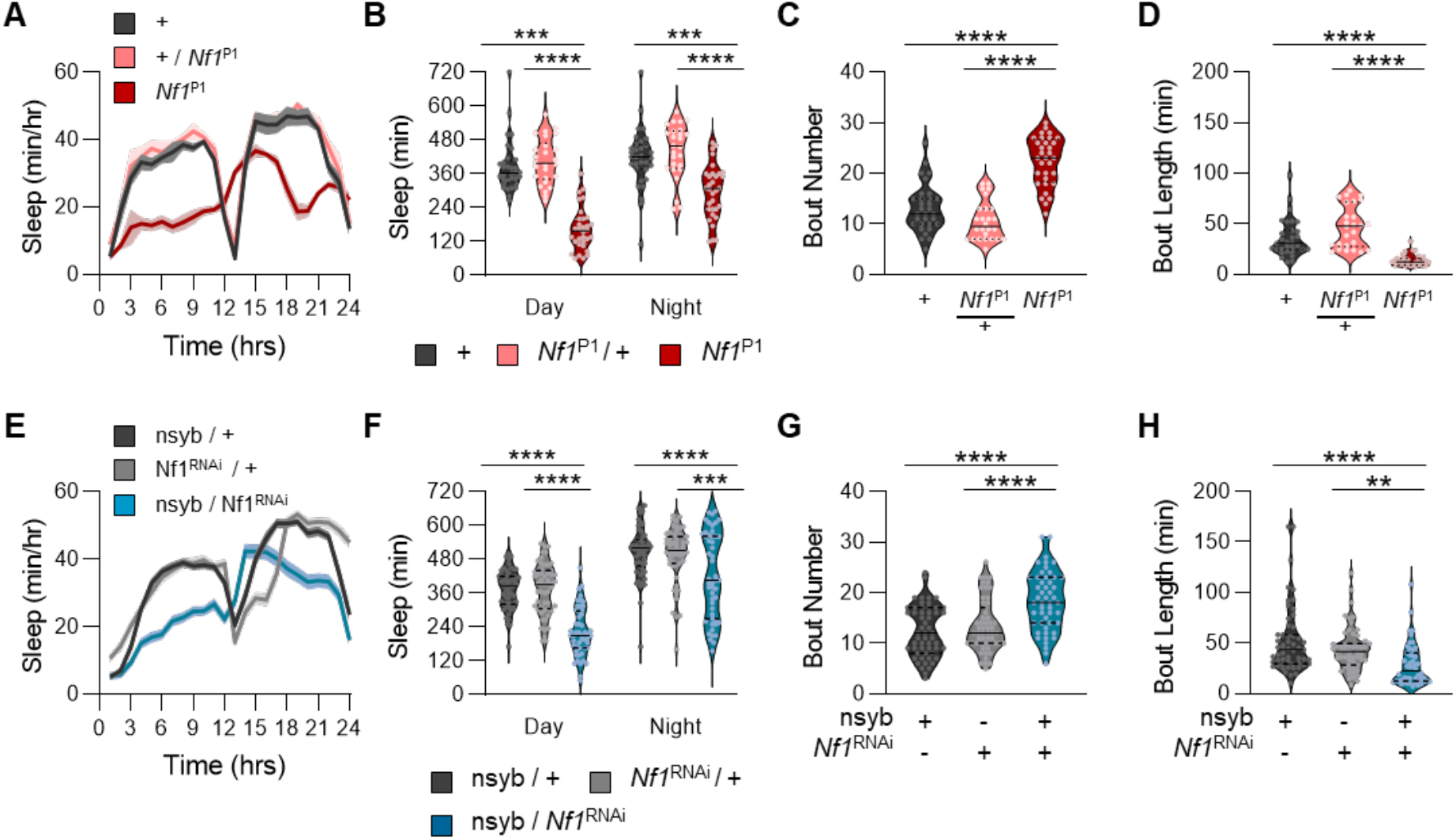
Loss of *Nf1* decreases sleep and increases sleep fragmentation. **(A-D)** Sleep traits of *Nf1*^P1^ mutants, heterozygotes, and their respective control. **A.** Sleep profiles. **B.** There is a significant effect of genotype on sleep duration (two-way ANOVA: F_2,174_ = 80.70, *P*<0.0001). Compared to control and heterozygote flies, *Nf1*^P1^ mutants sleep significantly less during the day (+, *P*<0.0001; het, *P*<0.0001) and night (+, *P*<0.0001; het, *P*<0.0001). **(C,D).** Compared to control and heterozygote flies, *Nf1*^P1^ mutants have a significantly higher **(C)** bout number (one-way ANOVA: F_2,86_ = 56.18, *P*<0.0001), and significantly lower **(D)** bout length (one-way ANOVA: F_2,86_ = 32.58, *P*<0.0001). **(E-H)** Sleep traits of pan-neuronal *Nf1*^RNAi^ knockdown flies and their respective controls. **E.** Sleep profiles. **F.** There is a significant effect of genotype on sleep duration (two-way ANOVA: F_2,314_ = 46.27, *P*<0.0001). Compared to controls, pan-neuronal knockdown of *Nf1* significantly reduces sleep during the day (nsyb>+, *P*<0.0001; *Nf1*^RNAi^>+, *P*<0.0001) and night (nsyb>+, *P*<0.0001; *Nf1*^RNAi^>+, *P*<0.0004). **(G,H)** Compared to controls, pan-neuronal knockdown of *Nf1* significantly increases **(G)** bout number (one-way ANOVA: F_2,157_= 18.35, *P*<0.0001), and significantly decreases **(H)** bout length (one-way ANOVA: F_2,157_ = 10.95, *P*<0.0001). For profiles, shaded regions indicate ± SEM. White background indicates daytime, while gray background indicates nighttime. ZT indicates zeitgeber time. For violin plots, the median (solid line), as well as the 25th and 75th percentiles (dotted lines) are shown. \*\**p*<0.01; \*\*\**p*<0.001; \*\*\*\**p*<0.0001.

To determine if the sleep and activity phenotypes of *Nf1* are due to loss of function in neurons, we selectively knocked down *Nf1* by expressing *Nf1*^RNAi^ under the control of the pan-neuronal driver nsyb-GAL4. Sleep was reduced and fragmented in flies upon pan-neuronal knockdown of *Nf1* in neurons (nsyb-GAL4>*Nf1*^RNAi^) compared to flies harboring either transgene alone (Fig 1E-H). Waking activity was also elevated with neuron-specific knockdown of *Nf1* (Fig S1B). To validate that these differences were not due to off-target effects of RNAi, we next confirmed these findings using an independently derived RNAi line (Zirin et al., 2020). We again found that pan-neuronal knockdown of *Nf1* significantly decreased sleep duration, while sleep fragmentation and waking activity increased significantly (Fig S2A-D). Therefore, pan-neuronal knockdown of *Nf1* fully recapitulates the mutant phenotype, suggesting that *Nf1* functions in neurons to regulate sleep.

To further investigate the role of *Nf1* on sleep consolidation, we analyzed activity patterns using a Markov model that predicts sleeping and waking propensity, indicators of sleep depth (Wiggin et al., 2020). In both *Nf1*^P1^ mutants and nsyb-GAL4>*Nf1*^RNAi^ flies, loss of *Nf1* increases the propensity to wake, while sleep propensity is reduced or remains unchanged (Fig S2E,F; Fig S3). Therefore, the phenotypes of both pan-neuronal RNAi knockdown of *Nf1* and genetic mutants further support the notion that *Nf1* promotes sleep and prevents sleep fragmentation.

Mounting evidence suggests that flies, like mammals, possess distinct sleep stages comprised of light and deep sleep (Faville et al., 2015; Alphen et al., 2021; Schafer and Keene, 2021). Increased arousal threshold, the phenomenon where a stronger stimulus is required to induce movement, is a key hallmark of sleep that is conserved across phyla (Alphen et al., 2021). Longer nighttime sleep bouts are associated with elevated arousal threshold, suggesting that sleep intensity increases during longer sleep bouts (Faville et al., 2015; Aplhen et al., 2021). To determine whether sleep depth is disrupted in *Nf1* deficient flies, we used the *Drosophila ARousal Tracking* (DART) system. We first implemented a paradigm that provides sleeping flies with increasing levels of vibration stimuli to determine the magnitude of the stimulus required to awaken the fly (Fig 2A; Faville et al., 2015). Arousal threshold was reduced during the day and the night in *Nf1*^P1^ mutant flies compared to wild-type controls and heterozygotes, revealing reduced sleep depth associated with loss of *Nf1* (Fig 2C). Similarly, arousal threshold was reduced during the day and night in flies with pan-neuronal knockdown of *Nf1* (nsyb-GAL4>*Nf1*^RNAI^) compared to flies harboring either transgene alone (Fig 2D). Sleep duration was also reduced in *Nf1*^P1^ mutants and upon pan-neuronal knockdown of *Nf1* in this system (Fig S4A,B). Together, these findings reveal that neuronal *Nf1* is required for normal arousal threshold.

**Figure 2.**
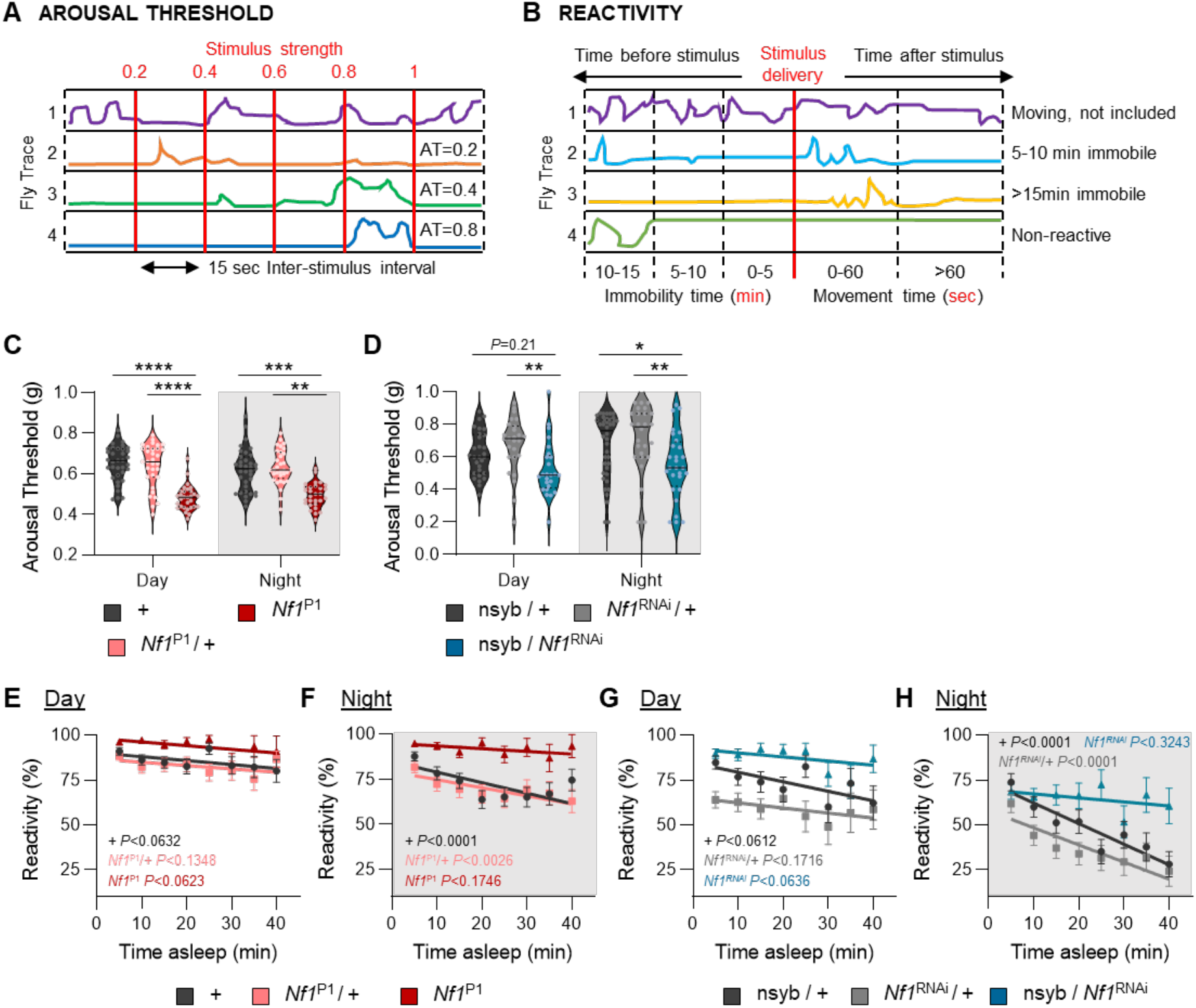
Loss of *Nf1* decreases sleep depth. **(A-G).** The *Drosophila* Arousal Tracking (DART) was used to probe arousal threshold and reactivity measurements. This system records fly movement while simultaneously controlling mechanical stimuli via a digital analog converter (DAC). All measurements included were taken from sleeping flies and determined hourly, starting at ZT0. **A.** Mechanical stimuli of increasing strength were used to assess arousal threshold and was determined by fly movement within 15 sec of stimulus delivery. **B.** Mechanical stimuli at maximum intensity was used to measure reactivity. A fly was considered reactive if it moved within 60 sec of stimulus delivery. **C.** There is a significant effect of genotype on arousal threshold (REML: F_2,94_ = 37.62, *P*<0.0001; N = 29–35). Compared to control and heterozygote flies, arousal threshold significantly decreases in *Nf1*^P1^ mutants and occurs during the day (+, *P*<0.0001; het, *P*<0.0001) and night (+, *P*<0.0001; het, *P*<0.0001). **D.** There is a significant effect of genotype on arousal threshold (REML: F_2,98_ = 7.795, *P*<0.0007; N = 25–40). Compared to controls, panneuronal knockdown of *Nf1* has mixed effects on arousal threshold during the day (nsyb>+, *P*<0.2138; *Nf1*^RNAi^>+, *P*<0.0040) and significantly decreases arousal threshold during the night (nsyb>+, *P*<0.0271; *Nf1* >+, *P*<0.00062). **E.** Linear regression of daytime reactivity as a function of time asleep in *Nf1*^P1^ mutants, heterozygotes, and their respective control. The slopes of each regression line are not significantly different from each other (ANCOVA with time asleep as the covariate: F_2,2062_ = 0.0254, *P*<0.9749). **F.** Linear regression of nighttime reactivity as a function of time asleep in *Nf1*^P1^ mutants, heterozygotes, and their respective control. The slopes of each regression line are significantly different from each other (F_2,2100_ = 57.05, *P*<0.0001). **G.** Linear regression of daytime reactivity as a function of time asleep in pan-neuronal *Nf1*^RNAi^ knockdown flies and their controls. The slopes of each regression line are not significantly different from each other (F_2,1285_ = 0.5551, *P*<0.5741). **H.** Linear regression of nighttime reactivity as a function of time asleep in pan-neuronal *Nf1*^RNAi^ knockdown flies and their controls. The slopes of each regression line are significantly different from each other (F_2,1564_ = 4.887, *P*<0.0077). For arousal threshold measurements, the median (solid line), as well as the 25th and 75th percentiles (dotted lines) are shown. For reactivity, error bars indicate ± SEM. The *P*-values in each panel indicates whether the slope of the regression line is significantly different from zero. White background indicates daytime, while gray background indicates nighttime. \**p*<0.05; \*\**p*<0.01; \*\*\**p*<0.001; \*\*\*\**p*<0.0001.

To determine whether *Nf1* is required for flies to modulate arousal threshold during individual sleep bouts we measured the reactivity of flies to a vibration stimulus and calculated their responsiveness as a function of time spent asleep prior to stimulus onset (Fig 2B). In control and heterozygous *Nf1*^P1^ flies, there was no effect of time spent asleep on reactivity during the day (Fig 2E). However, reactivity was significantly reduced during the night, as the slope of their respective regression lines differed significantly from zero (Fig 2F). *Nf1*^P1^ mutants had no effect on time spent asleep during the night or the day on reactivity (Fig 2E,F). Flies with pan-neuronal knockdown of *Nf1* maintained high levels of reactivity across sleep bouts of up to 40 minutes, phenocopying *Nf1*^P1^ mutants (Fig 2G,H). Therefore, *Nf1* is required for sleep duration-dependent changes in arousal threshold.

In both flies and mammals, sleep is associated with reduced metabolic rate (Brebbia & Altshuler, 1965; Brown et al., 2022; Caron & Stephenson, 2010; Katayose et al., 2009; Koban & Swinson, 2005; Stahl et al., 2017; White et al., 1985). To determine the effect of *Nf1* on sleep-dependent modulation of metabolic rate, we measured metabolic rate in awake and sleeping flies using the Sleep and Activity Metabolic Monitor (SAMM) system. This system uses indirect calorimetry to measure CO_2_ release, while simultaneously measuring activity via counting infrared beam crosses (Fig 3A; Brown et al., 2022; Stahl et al., 2017). In agreement with previous findings, sleep was reduced in *Nf1*^P1^ mutants in the SAMM system, and the total metabolic rate (VCO_2_) was elevated during the day and the night compared to controls (Fig S5A,B; Botero et al., 2021). Similar effects were observed upon pan-neuronal knockdown of *Nf1* (Fig S5C,D). To specifically examine the effects of sleep on CO_2_ output, we compared the overall CO_2_ output during waking and sleep. We found that CO_2_ output was significantly higher in *Nf1* mutant flies during both waking and sleeping, and was consistent during the day and night (Fig S6A,B). Pan-neuronal knockdown of *Nf1* (nsyb-GAL4>*Nf1*^RNAi^) similarly resulted in significantly higher CO_2_ output during both waking and sleeping (Fig S6C,D). This systematic dissection of CO_2_ output into sleep/waking states suggests that *Nf1* is required for the maintenance of metabolic rate.

**Figure 3.**
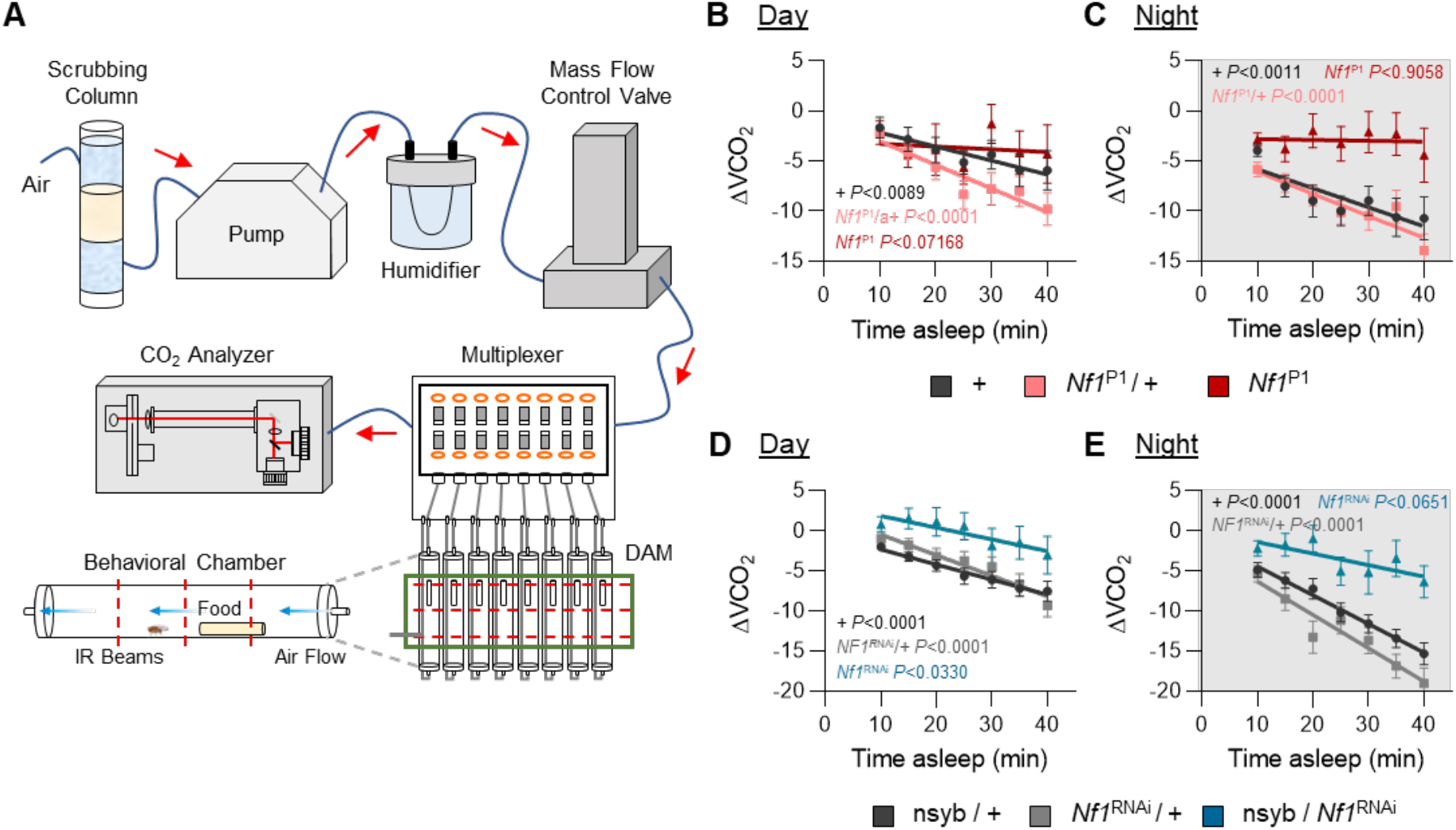
Loss of *Nf1* disrupts metabolic regulation of sleep. The Sleep and Metabolic Monitoring (SAMM) system was used to measure metabolic rate as a function of time spent asleep. **A.** Overview of the SAMM system. This system records activity while simultaneously measuring CO_2_ production, thereby enabling sleep and activity metrics to be paired with CO_2_ output. **B.** Linear regression of daytime CO_2_ output as a function of time asleep in *Nf1*^P1^ mutants, heterozygotes, and their respective control. There is no significant difference between the slopes of each regression line (ANCOVA with time asleep as the covariate: F_2,532_ = 2.988, *P*<0.0512). **C.** Linear regression of nighttime CO_2_ output as a function of time asleep in *Nf1* mutants, heterozygotes, and their respective control. The slopes of each regression line are significantly different from each other (F_2,538_ = 3.847, *P*<0.0219). **D.** Linear regression of daytime CO_2_ output as a function of time asleep in pan-neuronal *Nf1*^RNAi^ knockdown flies and their controls. There is no significant difference between the slopes of each regression line (F_2,805_ = 1.001, *P*<0.3680). **E.** Linear regression of nighttime CO_2_ output as a function of time asleep in pan-neuronal *Nf1*^RNAi^ knockdown flies and their controls. The slopes of each regression line are significantly different from each other (F_2,822_ = 5.625, *P*<0.0037). Error bars indicate ± SEM. The *P*-values in each panel indicates whether the slope of the regression line is significantly different from zero. White background indicates daytime, while gray background indicates nighttime. \**p*<0.05; \*\**p*<0.01; \*\*\*\**p*<0.0001.

To directly test whether sleep-metabolism interactions are disrupted by the loss of *Nf1*, we measured CO_2_ output over the length of a sleep bout. Metabolic rate was reduced during longer sleep bouts in control flies during the night, but did not change in *Nf1*^P1^ mutants, while there was no effect of sleep on metabolic rate during the day (Fig 3B,C). Similarly, pan-neuronal knockdown of *Nf1* (nsyb-GAL4>*Nf1*^RNAi^) abolished nighttime sleep-dependent changes in metabolic rate, while there was no effect of sleep on metabolic rate during the day (Fig 3D,E). These findings reveal a critical role for neuronal *Nf1* in sleep-dependent changes in metabolic rate. Taken together, loss of *Nf1* results in sleep fragmentation, reduced arousal threshold, and loss of sleep-dependent changes in metabolic rate, suggesting that *Nf1* is required for flies to enter deep sleep.

While *Nf1* is broadly expressed throughout the brain, its function has been linked to the modulation of GABA signaling during the formation of associative memories (Georganta et al., 2021). In *Drosophila*, the GABA_A_ receptor *Resistant to dieldrin (Rdl)* is expressed in numerous populations of sleep-regulating neurons (Fig 4A; Chung et al., 2009; Driscoll et al., 2021; Parisky et al., 2008). To examine whether *Nf1* functions in GABA_A_ receptor neurons, we selectively knocked down *Nf1* by expressing *Nf1*^RNAi^ under the control of *Rdl*-GAL4 and then measured its effect on sleep. Flies with *Nf1* knockdown in GABA_A_ receptor neurons (*Rdl*-GAL4>*Nf1*^RNAi^) slept less than control flies harboring either transgene alone (Fig 4B). Sleep was fragmented in *Rdl*-GAL4>*Nf1*^RNAi^ flies, with increased bout number, reduced bout length, and an increased propensity to wake (Fig 4C-D, Fig S7). Further, when sleep was measured in the DART and SAMM systems, knockdown of *Nf1* in GABA_A_ receptor neurons similarly reduced sleep duration (Fig S8A,B). Therefore, knockdown of *Nf1* in GABA_A_ receptor neurons phenocopies pan-neuronal knockdown of *Nf1*, suggesting that GABA-sensitive neurons contribute to the sleep abnormalities of *Nf1* mutant flies.

**Figure 4.**
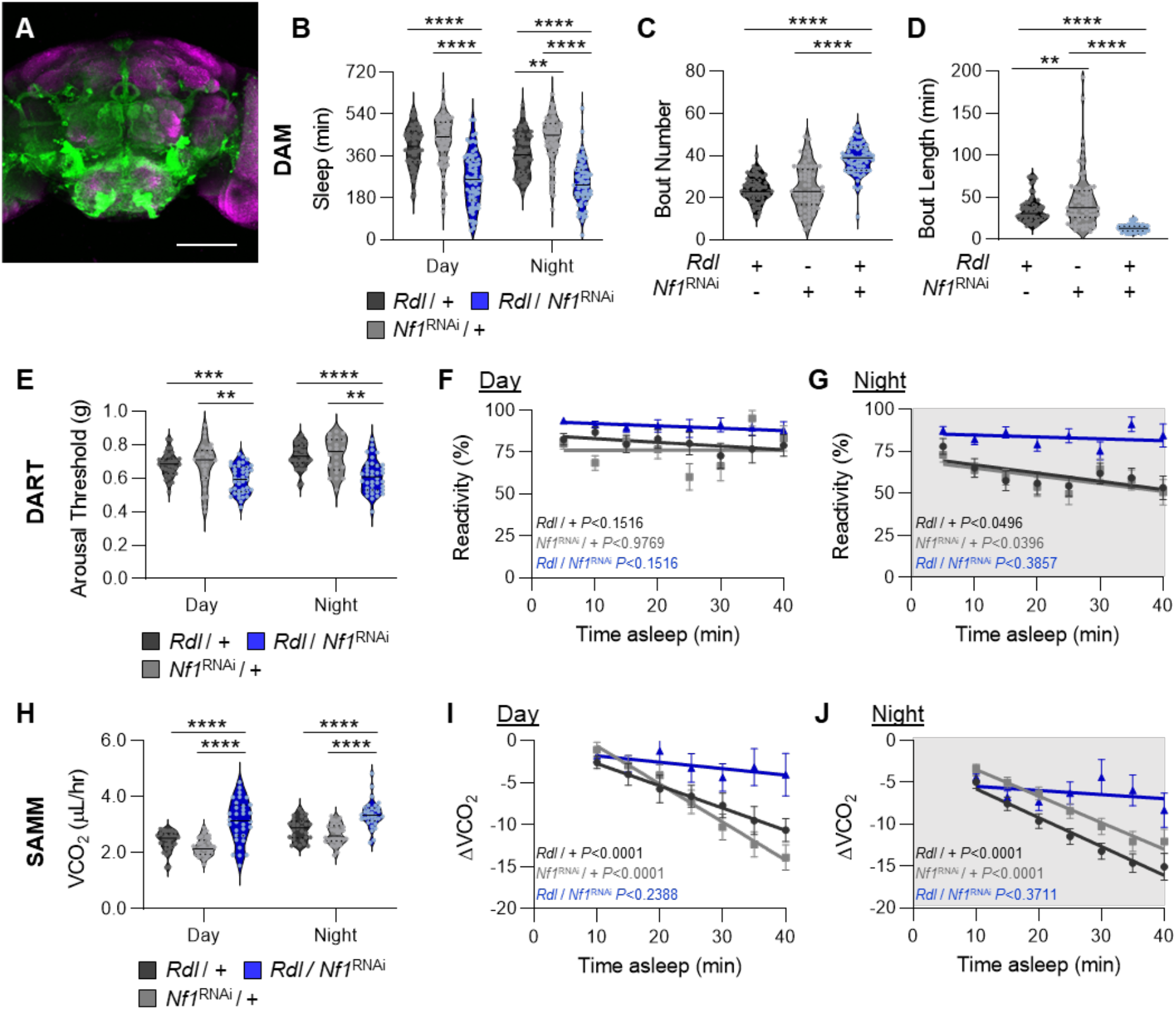
GABA_A_ receptor neurons mediate sleep depth via *Nf1*. GABA_A_ receptor neurons were targeted using the *Rdl*-GAL4 driver. **A.** The expression pattern of *Rdl*-expressing neurons is visualized with GFP. Background staining is NC82 antibody (magenta). Scale bar = 100μm. (B-D). Sleep traits of *Nf1*^RNAi^ knockdown flies and their respective controls. **B.** There is a significant effect of genotype on sleep duration (two-way ANOVA: F_2,386_ = 110.4, *P*<0.0001). Compared to controls, knockdown of *Nf1* in *Rdl*-expressing neurons significantly reduces sleep and occurs during the day (*Rdl*>+, *P*<0.0001; *Nf1*^RNAi^>+, *P*<0.0001) and night (*Rdl*>+, *P*<0.0001; *Nf1*^RNAi^ >+, *P*<0.0001). **(C,D)** Compared to controls, knockdown of *Nf1* in *Rdl*-expressing neurons significantly increases **(C)** bout number (one-way ANOVA: F_2,204_ = 9.007, *P*<0.0002), and significantly decreases **(D)** bout length (one-way ANOVA: F_2,204_= 47.04, *P*<0.0001). **(E-G)** Measurements of arousal threshold and reactivity in *Nf1*^RNAi^ knockdown flies and their respective controls using the DART system. **E.** There is a significant effect of genotype on arousal threshold (REML: F_2,103_ = 13.76, *P*<0.0001; N = 22–38). Compared to controls, knockdown of *Nf1* in *Rdl*-expressing neurons significantly decreases arousal threshold and occurs during the day (*Rdl*>+, *P*<0.0003; *Nf1*^RNAi^>+, *P*<0.0011) and night (*Rdl*>+, *P*<0.0001; *Nf1*^RNAi^>+, *P*<0.0014). **F.** Linear regression of daytime reactivity as a function of time asleep in *Nf1*^RNAi^ knockdown flies and their controls. The slopes of each regression line are not significantly different from each other (F_2,1769_ = 0.6085, *P*<0.5443). **G.** Linear regression of nighttime reactivity as a function of time asleep in *Nf1*^RNAi^ knockdown flies and their controls. The slopes of each regression line are significantly different from each other (F_2,1946_ = 2.804, *P*<0.0450). **(H-J)** Measurements of metabolic rate in *Nf1*^RNAi^ knockdown flies and their respective controls using the SAMM system. **H.** There is a significant effect of genotype on metabolic rate (two-way ANOVA: F_2,188_ = 55.60, *P*<0.0001). Compared to controls, knockdown of *Nf1* in *Rdl*-expressing neurons significantly increases CO_2_ output and occurs during the day (*Rdl*>+, *P*<0.0001; *Nf1*^RNAi^>+, *P*<0.0001) and night (*Rdl*>+, *P*<0.0001; *Nf1*^RNAi^>+, *P*<0.0001). **I.** Linear regression of daytime CO_2_ output as a function of time asleep in pan-neuronal *Nf1*^RNAi^ knockdown flies and their controls. The slopes of each regression line are significantly different from each other (F_2,698_ = 11.17, *P*<0.0001). **J.** Linear regression of nighttime CO_2_ output as a function of time asleep in pan-neuronal *Nf1*^RNAi^ knockdown flies and their controls. The slopes of each regression line are significantly different from each other (F_2,756_ = 13.64, *P*<0.0001). For violin plots, the median (solid line) as well as 25th and 75th percentiles (dotted lines) are shown. For reactivity and metabolic rate measurements, error bars indicate ± SEM. The *P*-values in each panel indicate whether the slope of the regression line is significantly different from zero. White background indicates daytime, while gray background indicates nighttime. \**p*<0.05; \*\**p*<0.01; \*\*\**p*<0.001; \*\*\*\**p*<0.0001.

It is possible that the effects on sleep duration and sleep depth are regulated by shared or distinct populations of neurons. Therefore, we sought to determine whether loss of *Nf1* in GABA_A_ receptor neurons also impacts arousal threshold and sleep-dependent changes in metabolic rate. Similar to pan-neuronal knockdown, arousal threshold was reduced in *Rdl*-GAL4> *Nf1*^RNAi^ flies, and these flies do not increase reactivity during long nighttime sleep bouts (Fig 4E-G). The metabolic phenotypes of *Nf1* mutant and pan-neuronal knockdown flies were also present in flies upon knockdown of *Nf1* in GABA_A_ receptor neurons. First, total metabolic rate was significantly increased, phenocopying pan-neuronal loss of *Nf1* (Fig 4H). Knockdown of *Nf1* in GABA_A_ receptor neurons similarly resulted in significantly higher CO_2_ output during both waking and sleeping states (Fig S8C,D). In addition, knockdown of *Nf1* in GABA_A_ receptor neurons abolished sleep-dependent changes in metabolic rate during the day and night (Fig 4I-J). Therefore, *Nf1* is required in GABA_A_ neurons to regulate sleep duration, arousal threshold, and sleep-dependent changes in metabolic rate.

In *Drosophila*, sleep loss is associated with shortened lifespan (Bushey et al., 2010; Koh et al., 2008; Vaccaro et al., 2020). To examine whether disrupted sleep impacts lifespan, we measured the effects of loss of *Nf1* on longevity in individually housed flies. Lifespan was significantly reduced in *Nf1*^P1^ mutant flies, as well as pan-neuronal knockdown (nsyb-GAL4>*Nf1*^RNAI^) or GABA_A_-receptor specific knockdown (*Rdl*-GAL4> *Nf1*^RNAI^) of *Nf1*, compared to their respective controls (Fig 5A, Fig S9A,B). These findings suggest that the loss of *Nf1* affects sleep duration and sleep quality and results in a significantly reduced lifespan.

**Figure 5.**
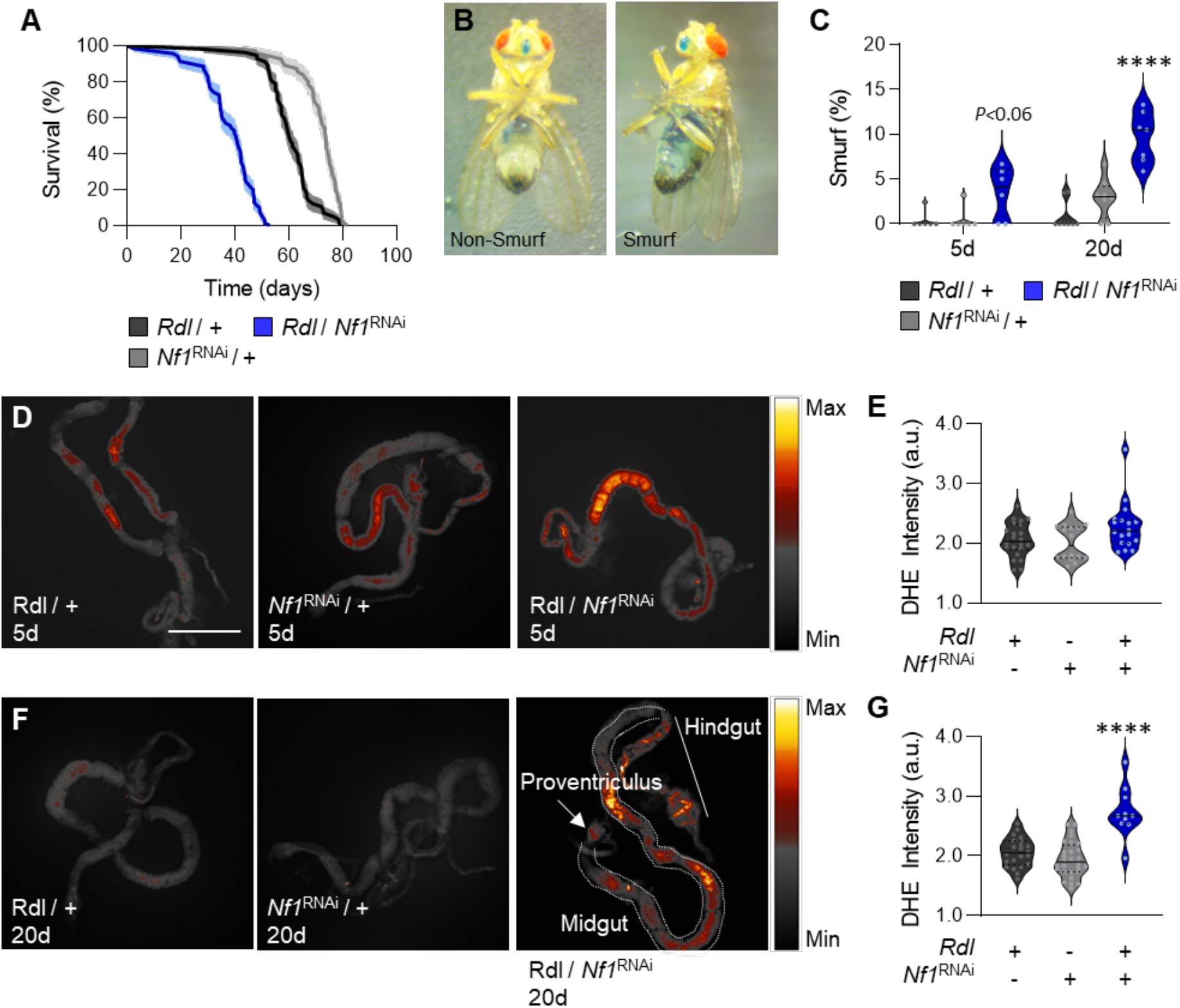
Loss of *Nf1* in GABA_A_ receptor neurons reduce longevity and promote aging-associated phenotypes. **A.** Compared to controls, knockdown of *Nf1* in *Rdl*-expressing neurons significantly decreases longevity (Log-Rank test: χ^2^ =253.4, d.f.=2, *P*<0.0001). **(B,C)** The Smurf assay was used to measure intestinal barrier dysfunction. **B.** Representative images depicting non-Smurf (left) and Smurf flies (right). **C.** There is a significant effect of genotype on intestinal permeability (two-way ANOVA: F_2,35_ = 29.45, *P*<0.0001). Knockdown of *Nf1* in *Rdl*-expressing neurons does not change intestinal barrier dysfunction in 5d flies (*Rdl*>+, *P*<0.0565; *Nf1*^RNAi^>+, *P*<0.0648), but significantly increases in 20d flies (*Rdl*>+, *P*<0.0001; *Nf1*^RNAi^>+, *P*<0.0001). **(D-F)** ROS was measured in 5d and 20d flies by quantifying oxidized DHE. Scale bar = 500μm. **D** Oxidized DHE was measured in 5d control and *Nf1* knockdown flies. **E.** There is no significant difference in oxidized DHE signal intensity in 5d flies (one-way ANOVA: F_2,53_ = 3.367, *P*<0.0520). **F.** Oxidized DHE was measured in 20d control and *Nf1* knockdown flies. **G.** Knockdown of *Nf1* in *Rdl*-expressing neurons significantly increases oxidized DHE signal intensity in 20d flies (one-way ANOVA: F_2,57_= 25.71, *P*<0.0001). The median (solid line) as well as 25th and 75th percentiles (dotted lines) are shown. \*\*\*\**p*<0.0001.

We next sought to measure the functional consequences of loss of *Nf1*. In *Drosophila* and mammals, chronic sleep loss is associated with deficiencies in gut homeostasis, that can result in death (Vaccaro et al., 2020). To measure gut integrity, flies were fed blue dye and then assayed for gut permeability (Martins et al., 2018; Rera et al., 2012). In control flies, gut permeability remains intact in young (5 days) and aged (20 days) flies (Fig 5B,C). However, in *Nf1*^P1^ mutants and flies with pan-neuronal knockdown of *Nf1*, gut permeability significantly increased in aged flies compared to controls (Fig S9C,D). Similarly, knockdown of *Nf1* in GABA_A_ receptor neurons (*Rdl*-GAL4>*Nf1*^RNAI^) significantly increased intestinal permeability in aged flies (Fig 5C). Together, these findings reveal that neuronal loss of *Nf1*, as well as selective loss in GABA_A_ receptor neurons, results in increased gut permeability that has been associated with aging.

It has previously been reported that sleep deprivation induces the generation of reactive oxygen species (ROS) that underlies reduced gut function and ultimately death (Vaccaro et al., 2020). These findings suggest that low sleep quality negatively impacts the health and longevity of *Drosophila*. To determine if loss of *Nf1* impairs gut function, we measured ROS levels in the gut in young (5 days) and aged (20 days) flies. Pan-neuronal knockdown of *Nf1* (nsyb-GAL4>*Nf1*^RNAI^) led to increased ROS levels in both young and aged flies compared to controls (Fig S10). Furthermore, ROS levels were also elevated in aged flies upon knockdown of *Nf1* in GABA_A_ receptor neurons (Fig 5D-G). Therefore, these findings support the notion that reduced gut function contributes to the reduced lifespan associated with the loss of *Nf1*.

## Discussion

Clinical evidence reveals Neurofibromin type 1 (*Nf1)* to be critical for regulating diverse biological functions, as humans afflicted with Neurofibromatosis Type 1 have behavioral manifestations including a high co-morbidity with ADHD, autism, learning impairments, and sleep disruption (Garg et al., 2013; Gutmann et al., 2017; Hyman et al., 2005, 2006; Leschziner et al., 2013; Licis et al., 2013; Walsh et al., 2013). Studies in mammalian models have revealed a robust role for *Nf1* in sleep and metabolic regulation, raising the possibility that they contribute to the complex systems in humans (Anastasaki et al., 2019; Tritz et al., 2021). In *Nf1*-deficient *Drosophila*, metabolic rate is chronically elevated, sleep is shortened, and circadian rhythms are dysregulated in *Nf1* mutants (Bai et al., 2018; Bai & Sehgal, 2015; Botero et al., 2021; King et al., 2016; Machado Almeida et al., 2021; Maurer et al., 2020; Williams et al., 2001), suggesting deep evolutionary conservation of *Nf1* function. The findings that *Nf1* plays a conserved role in regulating both sleep and metabolic rate raises the possibility that *Nf1* is a critical integrator of these processes and that it may play a role in various forms of metabolic dysfunction that are associated with sleep disturbance (Arble et al., 2015; Depner et al., 2014).

In *Drosophila* and mammals, sleep is associated with reduced metabolic rate (Brown et al., 2022; Fontvieille et al., 1994; Stahl et al., 2017). In *Drosophila*, metabolic rate is elevated across the circadian cycle in *Nf1* mutants, and this is associated with reductions in energy stores and starvation resistance (Botero et al., 2021). Here, we have applied indirect calorimetry to examine metabolic rate across individual sleep bouts and find that *Nf1* is required for reductions in metabolic rate associated with prolonged time spent asleep. These findings raise the possibility that *Nf1* signaling is a sleep output that specifically serves to regulate metabolic rate. Supporting this notion, *Nf1* is proposed to be an output of the circadian clock because circadian gene expression is normal in *Nf1* mutants, yet flies are arrhythmic (Williams et al., 2001), and *Nf1* impacts the physiology of neurons downstream of clock circuits (Bai et al., 2018). In *Drosophila*, numerous populations of neurons contribute to regulating sleep and wakefulness, presenting a challenge to localizing the integration of sleep and metabolic regulation (Shafer & Keene, 2021). The failure of *Nf1* mutant flies to integrate sleep and metabolic rate may provide a pathway to identify output from sleep neurons that regulate metabolic state.

There is growing evidence that *Drosophila*, like mammals, possess light and deep sleep (Faville et al., 2015). For example, readouts of both broad electrical activity and defined neural circuits suggest light sleep associates with periods early in a sleep bout, while deeper sleep associates with periods later in a sleep bout (Nitz et al., 2002; Tainton-Heap et al., 2021; van Alphen et al., 2013). Furthermore, periods later in a sleep bout are associated with an elevated arousal threshold that indicates deeper sleep (van Alphen et al., 2013). These findings are supported by functional evidence that consolidation of sleep bouts is required for critical brain functions, including waste clearance and memory, and that these processes are impaired when sleep is disrupted (Dissel et al., 2015; Tononi & Cirelli, 2014; van Alphen et al., 2021). Here, we provide evidence that *Nf1* flies fail to enter deep sleep, including sleep fragmentation, mathematical modeling of sleep pressure, reduced arousal threshold, and a loss of sleep-dependent reductions in metabolic rate. These findings suggest that *Nf1* mutants fail to enter deep sleep, even during prolonged sleep bouts. Examining local field potentials and neural activity within *Nf1* mutants, as has been previously described (Tainton-Heap et al., 2021; van Alphen et al., 2013), is likely to inform the neural basis for the loss of deep sleep.

*Nf1* is broadly expressed and regulates numerous behaviors and brain functions. For many behaviors, *Nf1* function has been localized to different subsets of neurons, suggesting localized changes in *Nf1* regulate distinct behaviors. For example, *Nf1* is broadly required within the pacemaker circuit to regulate 24-hour rhythms, while *Nf1* in the mushroom bodies regulates clock-dependent wakefulness (Bai et al., 2018; Machado Almeida et al., 2021). For a number of other processes, including grooming behavior and metabolic rate, the specific population of neurons where *Nf1* functions has not been identified (Botero et al., 2021; King et al., 2016). The broad and diverse effect of *Nf1* raises the possibility that *Nf1* functions widely in many circuits, and that it may be challenging to localize its function to defined cell types. Here, we find that the metabolic and sleep phenotypes of *Nf1* mutant flies are phenocopied in flies with specific loss of *Nf1* in GABA_A_ receptor/*Rdl*-expressing neurons. These findings raise the possibility that sleep dysregulation is due to altered GABA signaling. Supporting this notion, GABA signaling to the mushroom bodies, the *Drosophila* memory center, is dysregulated in *Nf1* mutants (Georganta et al., 2021). It has also previously been reported that *Nf1* knockdown in GABA_A_ receptor neurons leads to shortened sleep bouts and reduced sleep duration (Maurer et al., 2020). These findings support the notion that *Nf1* modulates GABA signaling. Future studies defining the specific population(s) of neurons where *Nf1* functions may reveal novel neural mechanisms regulating sleep-dependent regulation of metabolic rate.

Epidemiological data and individuals with chronic sleep loss reveal a link between shortened sleep duration and serious health problems (Chattu et al., 2018; Medic et al., 2017). Additionally, in several model organisms, sleep restriction can lead to premature death (Bentivoglio & Grassi-Zucconi, 1997; Koh et al., 2008; Rechtschaffen et al., 1983; Shaw et al., 2002; Stephenson et al., 2007). Lifespan is reduced in *Nf1* mutant flies (Tong et al., 2007), but its relationship to reduced sleep or circadian dysregulation has been unclear. In *Drosophila*, lifespan is reduced in short-sleeping genetic mutants and by chronic sleep deprivation (Bushey et al., 2010; Koh et al., 2008). Sleep loss induced by acute manipulations in young flies, or during aging, results in increased sensitivity to ROS, suggesting the generation of ROS, or changes in clearing ROS, may be a critical function of sleep that is necessary for survival (Hill et al., 2018; Koh et al., 2006; Wang et al., 2010). Further, evidence suggests that sleep deprivation leads to increased accumulation of ROS in the gut, resulting in gut permeability and death (Vaccaro et al., 2020). We find the ROS and permeability phenotypes in the guts of aged *Nf1* flies phenocopy those of animals that have been mechanically sleep deprived. These findings suggest that sleep consolidation, or deep sleep, is essential for maintaining fly health. Ultimately, genetic models with reduced sleep quality may resemble human sleep disorders more closely than chronic sleep deprivation.

Taken together, our findings reveal a novel and complex role for *Nf1* in regulating sleep depth. Loss of *Nf1* induces multiple phenotypes classically associated with a loss of deep sleep. Human mutations in *Nf1* have been introduced to *Drosophila* and phenocopy many aspects of the human disease (Botero et al., 2021; Hannan et al., 2006; Ho et al., 2007; Walker et al., 2006). Therefore, these findings establish *Nf1* mutants as a model to study the function of deep sleep and provide the ability to investigate the function of disease-causing mutations on sleep regulation.

## Methods

### Fly husbandry and Stocks

Flies were grown and maintained on standard *Drosophila* food media (Bloomington Recipe, Genesee Scientific, San Diego, California) in incubators (Powers Scientific, Warminster, Pennsylvania) at 25°C on a 12:12 LD cycle with humidity set to 55–65%. The following fly strains were obtained from the Bloomington Stock Center: *w*^1118^ (#5905); Nsyb-GAL4 (#39171; Jenett et al., 2012); *Rdl*-GAL4 (#66509); UAS-mcd8::GFP (#32186; Pfeiffer et al., 2010); and UAS-*Nf1*^RNAi2^ (#25845; Perkins et al., 2015). The *Nf1*^P1^ and UAS-dicer2;UAS-*Nf1*^RNAi^ lines were a kind gift from Seth Tomchik (Botero et al., 2021; Dietzl et al., 2007). All lines were backcrossed to the *w*^1118^ laboratory strain for 10 generations. Unless otherwise stated, 3-to-5 day old mated males were used for all experiments performed in this study. For experiments using aged flies, flies were maintained on standard food and transferred to fresh vials every other day.

### Sleep and activity

For experiments using the *Drosophila* Activity Monitoring (DAM) system (Trikinetics, Waltham, MA, USA), measurements of sleep and waking activity were measured as previously described (Hendricks et al., 2000; Shaw et al., 2000). For each individual fly, the DAM system measures activity by counting the number of infrared beam crossings over time. These activity data were then used to calculate sleep, defined as bouts of immobility of 5 min or more. Sleep traits were then extracted using the *Drosophila* Sleep Counting Macro (Pfeiffenberger et al., 2010).

### Arousal threshold and reactivity

Arousal threshold was measured using the *Drosophila* Arousal Tracking system (DART), as previously described (Faville et al., 2015). In brief, individual female flies were loaded into plastic tubes (Trikinectics, Waltham, Massachusetts) and placed onto trays containing vibrating motors. Flies were recorded continuously using a USB-webcam (QuickCam Pro 900, Logitech, Lausanne, Switzerland) with a resolution of 960×720 at 5 frames per second. The vibrational stimulus, video tracking parameters, and data analysis were performed using the DART interface developed in MATLAB (MathWorks, Natick, Massachusetts). To track fly movement, raw video flies were subsampled to 1 frame per second. Fly movement, or a difference in pixelation from one frame to the next, was detected by subtracting a background image from the current frame. The background image was generated as the average of 20 randomly selected frames from a given video. Fly activity was measured as movement of greater than 3 mm. Sleep was determined by the absolute location of each fly and was measured as bouts of immobility for 5 min or more. Arousal threshold was assessed using sequentially increasing vibration intensities, from 0 to 1.2 g, in 0.3 g increments, with an inter-stimulus delay of 15 s, once per hour over 24 hours starting at ZT0. Measurements of arousal threshold are reported as the proportion of the maximum force applied to the platform, thus an arousal threshold of 40% is 40% of 1.2g.

### Indirect calorimetry

Metabolic rate was measured using the Sleep and Activity Metabolic Monitor (SAMM) system, as previously described (Brown et al., 2022; Stahl et al., 2017). Briefly, male flies were placed individually into behavioral chambers containing a food vial of 1% agar and 5% sucrose. Flies were acclimated to the chambers for 24 hrs and then metabolic rate was assessed by quantifying the amount of CO_2_ produced in 5 min intervals during the subsequent 24hrs. To investigate how CO_2_ production may change with time spent asleep, sleep and activity were measured simultaneously using the *Drosophila* Locomotor Activity Monitor System. The percent change in VCO_2_ over the duration of a single sleep bout was calculated using the following equation: [(VCO_2_ @ 5 min) – (VCO_2_ @ 10 min)] / (VCO_2_ @ 5min) × 100. This was repeated for each 5 min bin of sleep, for the entire length of the sleep bout. Since a single fly typically has multiple sleep bouts, the percent change in VCO_2_ for each 5 min bin of sleep was averaged across all sleep bouts over the course of the day/night.

### Immunohistochemistry

Brains of three to five day old female flies were dissected in ice-cold phosphate buffered saline (PBS) and fixed in 4% formaldehyde, PBS, and 0.5% Triton-X for 35 min at room temperature, as previously described (Kubrak et al., 2016). Brains were then rinsed 3x with cold PBS and 0.5% Triton-X (PBST) for 10 min at room temperature and then overnight at 4°C. The following day, the brains were incubated for 24 hours in primary antibody (1:20 mouse nc82; Iowa Hybridoma Bank; The Developmental Studies Hybridoma Bank, Iowa City, Iowa), and then diluted in 0.5% PBST at 4°C on a rotator. The following day, the brains were rinsed 3x in cold PBST for 10 min at room temperature and then incubated in secondary antibody (1:400 donkey anti-rabbit Alexa 488 and 1:400 donkey anti-mouse Alexa 647; ThermoFisher Scientific, Waltham, Massachusetts) for 95 min at room temperature. The brains were again rinsed 3x in cold PBST for 10 min at room temperature, then stored overnight in 0.5% PBST at 4°C. Lastly, the brains were mounted in Vectashield (VECTOR Laboratories, Burlingame, California) between a glass slide and coverslip, and then imaged in 2μm sections on a Nikon A1R confocal microscope (Nikon, Tokyo, Japan) using a 20X oil immersion objective. Images are presented as the Z-stack projection through the entire brain and processed using ImageJ2.

### Longevity

Longevity was measured using the DAM system. Freshly emerged flies were isolated and provided time to mate for 2 days. Male flies were then separated by anesthetizing with mild CO_2_ and loaded into tubes containing standard food. Flies were flipped to new tubes containing fresh standard food every 5 days. The time of death was manually determined for each individual fly as the last bout of waking activity. The lifespan of a fly was calculated as the number of days it survived post-emergence.

### Intestinal permeability

Intestinal integrity was assessed using the Smurf assay, as previously described (Martins et al., 2018; Rera et al., 2012). First, freshly emerged flies were isolated and provided time to mate for 2 days. Male flies were then separated by anesthetizing with mild CO_2_ and placed into vials containing standard food at a density of ~20 flies per vial. At ZT 0, flies of a given age and genotype were transferred onto fresh medium containing blue dye (2.5% w/v; FD&C blue dye #1) for 24 hrs. At ZT 0 the following day, the percentage of Smurf flies in each vial was recorded. Flies were considered Smurf if blue coloration extended beyond the gut.

### ROS Imaging and Quantification

*In situ* ROS detection was performed using dihydroethidium (DHE; D11347, ThermoFisher Scientific), as previously described (Owusu-Ansah et al., 2008; Vaccaro et al., 2020). Briefly, flies were anesthetized on ice and whole guts were dissected in Gibco™ Schneider’s *Drosophila* Medium (21720024, ThermoFisher Scientific). The tissue was then incubated at room temperature with 60 μm DHE for 5 min in the dark. Next, tissues were washed 3x in Schneider’s medium for 5 min and then once in PBS for 5 min. Samples were then mounted in Vectashield Antifade Mounting Medium with DAPI (VECTOR Laboratories) between a glass slide and coverslip and then imaged immediately on a Nikon A1R confocal microscope (Nikon) using a 10X air objective. Total ROS levels were quantified from pixel intensities of the Z-stack projection (sum slices). An ROI (gut tissue) was determined from the DAPI channel and then the mean of the summed DHE intensity averaged from each tissue was used for statistical analysis. Images are presented as the Z-stack projection through the entire gut and were processed using ImageJ2.

### Statistical Analysis

Measurements of sleep duration, metabolic rate, and DHE intensity are presented as bar graphs displaying the mean ± standard error. Unless otherwise noted, a one-way or two-way analysis of variance (ANOVA) was used for comparisons between two or more genotypes and one treatment or two or more genotypes and two treatments, respectively. Measurements of arousal threshold and intestinal permeability were not normally distributed and so are presented as violin plots; indicating the median, 25^th^, and 75^th^ percentiles. The non-parametric Kruskal-Wallis test was used to compare two or more genotypes. To compare two or more genotypes and two treatments, a restricted maximum likelihood (REML) estimation was used. Linear regression analyses were used to characterize the relationship between the change in CO_2_ output and time spent asleep as well as between reactivity and time spent asleep. An F-test was used to determine whether the slope of each regression line was different from zero, while an ANCOVA was used to compare the slopes of different treatments. To assess differences in survivorship, longevity was analyzed using a log-rank test. All *post hoc* analyses were performed using Sidak’s multiple comparisons test. All statistical analyses were performed using InStat software (GraphPad Software 8.0).

## Acknowledgments

We are thankful to members of the Keene laboratory for helpful discussions and technical support. This work was supported by the National Institutes of Health [grant numbers: R21NS124198 to A.C.K and S.T, R01DC017390 to A.C.K, and K99AG071833 to E.B.B].

---

**Supplemental Figure 1. Loss of *Nf1* increases waking activity**. Waking activity was measured as the number of beam crosses per waking minute. **A**. Compared to control and heterozygote flies, *Nf1*^P1^ mutants have significantly higher waking activity (one-way ANOVA: F_2,86_ = 41.50, *P*<0.0001). **B**. Compared to controls, pan-neuronal knockdown of *Nf1* significantly increases waking activity (one-way ANOVA: F_2,157_ = 11.48, *P*<0.0001). The median (solid line) as well as 25th and 75th percentiles (dotted lines) are shown. \**p*<0.05; \*\*\*\**p*<0.0001.

**Supplemental Figure 2. Pan-neuronal knockdown of *Nf1* significantly reduces sleep duration and sleep depth using an independent RNAi line. (A-F).** Sleep and activity traits of pan-neuronal *Nf1*^RNAi2^ knockdown flies and their respective controls. **A**. There is a significant effect of genotype on sleep duration (two-way ANOVA: F_2,378_ = 79.47, *P*<0.0001). Compared to controls, pan-neuronal knockdown of *Nf1* significantly reduces sleep during the day (nsyb>+, *P*<0.0001; *Nf1*^RNAi2^>+, *P*<0.0001) and night (nsyb>+, *P*<0.0001; *Nf1*^RNAi2^>+, *P*<0.0001). **(B,C)** Compared to controls, pan-neuronal knockdown of *Nf1* has no effect on (**B**) bout number (one-way ANOVA: F_2,189_ = 2.422, *P*<0.0915), while (**C**) bout length is significantly lower (one-way ANOVA: F_2,189_ = 11.21, *P*<0.0001). **D**. Compared to controls, pan-neuronal knockdown of *Nf1* significantly increases waking activity (one-way ANOVA: F_2,189_ = 18.37, *P*<0.0001). **E**. P(Doze) is significantly lower upon knockdown of *Nf1* (one-way ANOVA: F_2,189_ = 60.44, *P*<0.0001). **F**. P(Wake) is significantly higer upon knockdown of *Nf1* (one-way ANOVA: F_2,189_ = 33.11, *P*<0.0001). **(G,H)** Linear regression of **(G)** daytime and **(H)** nighttime reactivity as a function of time asleep in *Nf1*^RNAi^ knockdown flies and their controls. During the day, the slopes of each regression line are not significantly different from each other (F_2,816_ = 1.243, *P*=0.2890). During the night, the slopes of each regression line are significantly different from each other (F_2,823_ = 3.504, *P*=0.0305). For violin plots, the median (solid line) as well as 25th and 75th percentiles (dotted lines) are shown. For reactivity measurements, error bars indicate ± SEM. The *P*-values in each panel indicate whether the slope of the regression line is significantly different from zero. White background indicates daytime, while gray background indicates nighttime. \*\**p*<0.01; \*\*\**p*<0.001; \*\*\*\**p*<0.0001.

**Supplemental Figure 3. Probabilistic analysis suggests that loss of *Nf1* increases the probability of waking. (A-D)** Computational modeling of waking probabilities. **A**. Profiles of the probability of waking up in *Nf1*^P1^ mutants, heterozygotes, and their control. **B**. There is a significant effect of genotype on the probability of waking up (two-way ANOVA: F_2,172_ = 144.6, *P*<0.0001). P(Wake) is significantly higher in *Nf1*^P1^ mutant flies during the day (+, *P*<0.0001; het, *P*<0.0001) and night (+, *P*<0.0001; het, *P*<0.0001). **C**. Profiles of the probability of waking up in pan-neuronal *Nf1*^RNAi^ knockdown flies and their controls. **D**. There is a significant effect of genotype on the probability of waking up (two-way ANOVA: F_2,314_ = 67.34, *P*<0.0001). P(Wake) is significantly higher upon knockdown of *Nf1* during the day (nsyb>+, *P*<0.0001; *Nf1*^RNAi^>+, *P*<0.0001) and night (nsyb>+, *P*<0.0001; *Nf1*^RNAi^>+, *P*<0.0001). **(E-H)** Computational modeling of sleep probabilities. **E**. Profiles of the probability of falling asleep in *Nf1*^P1^ mutants, heterozygotes, and their control. **F**. There is a significant effect of genotype on the probability of falling asleep (two-way ANOVA: F_2,172_ = 23.10, *P*<0.0001). P(Doze) is significantly lower in *Nf1*^P1^ mutant flies during the day (+, *P*<0.0001; het, *P*<0.0001), but there is no difference during the night (+, *P*<0.0001; het, *P*<0.0001). **G**. Profiles of the probability of falling asleep in pan-neuronal *Nf1*^RNAi^ knockdown flies and their controls. **H**. There is a significant effect of genotype on the probability of falling asleep (two-way ANOVA: F_2,314_ = 13.55, *P*<0.0001). P(Doze) is significantly lower upon knockdown of *Nf1* during the day (nsyb>+, *P*<0.0001; *Nf1*^RNAi^>+, *P*<0.0001), but there is no difference during the night (nsyb>+, *P*<0.8053; *Nf1*^RNAi^>+, *P*<0.9999). For profiles, shaded regions indicate ± SEM. White background indicates daytime, while gray background indicates nighttime. ZT indicates zeitgeber time. For violin plots, the median (solid line) as well as 25th and 75th percentiles (dotted lines) are shown. \*\*\*\**p*<0.0001.

**Supplemental Figure 4. Loss of *Nf1* decreases sleep in the DART system**. **A**. There is a significant effect of genotype on sleep duration (two-way ANOVA: F_2,372_ = 131.7, *P*<0.0001). Compared to control and heterozygote flies, *Nf1*^P1^ mutants sleep significantly less during the day (+, *P*<0.0001; het, *P*<0.0001) and night (+, *P*<0.0001; het, *P*<0.0001). **B**. There is a significant effect of genotype on sleep duration (two-way ANOVA: F_2,220_ = 19.05, *P*<0.0001). Compared to controls, pan-neuronal knockdown of *Nf1* significantly reduces sleep during the day (nsyb>+, *P*<0.0003; *Nf1*^RNAi^>+, *P*<0.0001) and night (nsyb>+, *P*<0.0106; *Nf1*^RNAi^>+, *P*<0.0135). The median (solid line) as well as 25th and 75th percentiles (dotted lines) are shown. \**p*<0.05; \*\*\**p*<0.001; \*\*\*\**p*<0.0001.

**Supplemental Figure 5. Loss of *Nf1* decreases sleep and increases metabolic rate in the SAMM system**. Sleep duration and metabolic rate were measured in the SAMM system. **A**. There is a significant effect of genotype on sleep duration (two-way ANOVA: F_2,154_ = 10.92, *P*<0.0001). Compared to control and heterozygote flies, *Nf1*^P1^ mutants sleep significantly less during the day (+, *P*<0.0414; het, *P*<0.0159) and night (+, *P*<0.0167; het, *P*<0.0033). **B**. There is a significant effect of genotype on metabolic rate (two-way ANOVA: F_2,154_ = 43.72, *P*<0.0001). Compared to control and heterozygote flies, *Nf1*^P1^ mutants significantly increase CO_2_ output during the day (+, *P*<0.0001; het, *P*<0.0001) and night (+, *P*<0.0001; het, *P*<0.0001). **C**. There is a significant effect of genotype on sleep duration (two-way ANOVA: F_2,224_ = 13.79, *P*<0.0001). Compared to controls, pan-neuronal knockdown of *Nf1* significantly decreases sleep during the day (nsyb>+, *P*<0.0001; *Nf1*^RNAi^>+, *P*<0.0001), but not the night (nsyb>+, *P*<0.3333; *Nf1*^RNAi^>+, *P*<0.2203). **D**. There is a significant effect of genotype on metabolic rate (two-way ANOVA: F_2,224_ = 136.0, *P*<0.0001). Compared to controls, pan-neuronal knockdown of *Nf1* significantly increases CO_2_ output during the day (nsyb>+, *P*<0.0001; *Nf1*^RNAi^ >+, *P*<0.0001) and night (nsyb>+, *P*<0.0001; *Nf1*^RNAi^ >+, *P*<0.0001). The median (solid line) as well as 25th and 75th percentiles (dotted lines) are shown. \**p*<0.05; \*\**p*<0.01; \*\*\*\**p*<0.0001.

**Supplemental Figure 6. Loss of *Nf1* increases metabolic rate during waking and sleeping**. **A**. There is a significant effect of genotype on metabolic rate during waking (two-way ANOVA: F_2,154_ = 16.76, *P*<0.0001). In the daytime, *Nf1*^P1^ mutants significantly increase waking CO_2_ output compared to control and heterozygote flies (+, *P*<0.0003; het, *P*<0.0412). At night, *Nf1*^P1^ mutants significantly increase waking CO_2_ output compared to control flies (+, *P*<0.0002), with heterozygotes being intermediate (het, *P*<0.0762). **B**. There is a significant effect of genotype on metabolic rate during sleep (two-way ANOVA: F_2,154_ = 53.48, *P*<0.0001). Compared to control and heterozygote flies, *Nf1*^P1^ mutants significantly increase CO_2_ output during sleep during the day (+, *P*<0.0001; het, *P*<0.0001) and night (+, *P*<0.0001; het, *P*<0.0001). **C**. There is a significant effect of genotype on metabolic rate during waking (two-way ANOVA: F_2,224_ = 97.10, *P*<0.0001). Compared to controls, pan-neuronal knockdown of *Nf1* significantly increases waking CO_2_ output during the day (nsyb>+, *P*<0.0001; *Nf1*^RNAi^>+, *P*<0.0001) and night (nsyb>+, *P*<0.0001; *Nf1*^RNAi^>+, *P*<0.0001). **D**. There is a significant effect of genotype on metabolic rate during sleep (two-way ANOVA: F_2,224_ = 176.1, *P*<0.0001). Compared to controls, pan-neuronal knockdown of *Nf1* significantly increases CO_2_ output during sleep during the day (nsyb>+, *P*<0.0001; *Nf1*^RNAi^>+, *P*<0.0001) and night (nsyb>+, *P*<0.0001; *Nf1*^RNAi^>+, *P*<0.0001). The median (solid line) as well as 25th and 75th percentiles (dotted lines) are shown. \**p*<0.05; \*\*\**p*<0.001; \*\*\*\**p*<0.0001.

**Supplemental Figure 7. Computational modeling of sleep and waking probabilities upon knockdown of *Nf1* in *Rdl*-expressing neurons. A**. Compared to controls, P(Dose) is significantly lower upon knockdown of *Nf1* in *Rdl*-expressing neurons (one-way ANOVA: F_2,193_ = 3.865, *P*<0.0226). **B**. Compared to controls, P(Wake) is significantly higher upon knockdown of *Nf1* in *Rdl*-expressing neurons (one-way ANOVA: F_2,193_ = 8.317, *P*<0.0003). The median (solid line) as well as 25th and 75th percentiles (dotted lines) are shown. \*\**p*<0.01; \*\*\**p*<0.001; \*\*\*\**p*<0.0001.

**Supplemental Figure 8. Knockdown of *Nf1* in *Rdl*-expressing neurons decreases sleep and increases metabolic rate**. **A**. There is a significant effect of genotype on sleep duration in the DART system (two-way ANOVA: F_2,378_ = 27.38, *P*<0.0001). Compared to controls, knockdown of *Nf1* in *Rdl*-expressing neurons significantly reduces sleep and occurs during day (*Rdl*>+, *P*<0.0012; *Nf1*^RNAi^>+, *P*<0.0001) and night (*Rdl*>+, *P*<0.0015; *Nf1*^RNAi^>+, *P*<0.0001). **B**. There is a significant effect of genotype on sleep duration in the SAMM system (two-way ANOVA: F_2,188_ = 4.708, *P*<0.0101). Compared to controls, knockdown of *Nf1* in *Rdl*-expressing neurons significantly reduces sleep, but only occurs during the day (*Rdl*>+, *P*<0.0001; *Nf1*^RNAi^>+, *P*<0.0001) and not the night (*Rdl*>+, *P*<0.9410; *Nf1*^RNAi^>+, *P*<0.2371). **C**. There is a significant effect of genotype on metabolic rate during waking (two-way ANOVA: F_2,188_ = 54.14, *P*<0.0001). Compared to controls, knockdown of *Nf1* in *Rdl*-expressing neurons significantly increases CO_2_ output during the day (*Rdl*>+, *P*<0.0001; *Nf1*^RNAi^>+, *P*<0.0001) and night (*Rdl*>+, *P*<0.0003; *Nf1*^RNAi^>+, *P*<0.0001). **D**. There is a significant effect of genotype on metabolic rate during sleep (two-way ANOVA: F_2,188_ = 136.1, *P*<0.0001). Compared to controls, knockdown of *Nf1* in *Rdl*-expressing neurons significantly increases CO_2_ output during the day (*Rdl*>+, *P*<0.0001; *Nf1*^RNAi^>+, *P*<0.0001) and night (*Rdl*>+, *P*<0.0001; *Nf1*^RNAi^>+, *P*<0.0001). The median (solid line) as well as 25th and 75th percentiles (dotted lines) are shown. \*\**p*<0.01; \*\*\*\**p*<0.0001.

**Supplemental Figure 9. Loss of *Nf1* promotes aging-associated phenotypes. A**. Compared to control flies, loss of *Nf1* significantly decreases longevity (Log-Rank test: χ^2^=209.0, d.f.=2, *P*<0.0001). **B**. Compared to controls, pan-neuronal knockdown of *Nf1* significantly decreases longevity (Log-Rank test: χ^2^=253.4, d.f.=2, *P*<0.0001). **C.** There is a significant effect of genotype on intestinal permeability (two-way ANOVA: F1,42 = 29.45, *P*<0.0002). Loss of *Nf1* does not change intestinal barrier dysfunction in 5d flies (+, *P*<0.0565; het, *P*<0.0648), but significantly increases in 20d flies (+, *P*<0.0001; het, *P*<0.0001). **D.** There is a significant effect of genotype on intestinal permeability (two-way ANOVA: F_2,65_ = 18.80, *P*<0.0001). Pan-neuronal knockdown of *Nf1* does not change intestinal barrier dysfunction in 5d flies (nsyb>+, *P*<0.2093; *Nf1*^RNAi^>+, *P*<0.1973), but significantly increases in 20d flies (nsyb>+, *P*<0.0001; *Nf1*^RNAi^>+, *P*<0.0001). The median (solid line) as well as 25th and 75th percentiles (dotted lines) are shown. \*\*\**p*<0.001; \*\*\*\**p*<0.0001.

**Supplemental Figure 10. ROS in the gut increases upon pan-neuronal knockdown of *Nf1*.** ROS was measured in 5d and 20d flies by quantifying oxidized DHE levels. **A.** Oxidized DHE was measured in 5d control and *Nf1* knockdown flies. **B**. Pan-neuronal knockdown of *Nf1* significantly increases oxidized DHE signal intensity in 5d flies (one-way ANOVA: F_2,51_ = 30.20, *P*<0.0001). **C**. Oxidized DHE was measured in 20d control and *Nf1* knockdown flies. **D**. Pan-neuronal knockdown of *Nf1* significantly increases oxidized DHE signal intensity in 20d flies (one-way ANOVA: F_2,47_ = 16.40, *P*<0.0001). Scale bar = 500μm. The median (solid line) as well as 25th and 75th percentiles (dotted lines) are shown. \*\*\**p*<0.001; \*\*\*\**p*<0.0001.

## Notes

### Competing Interest Statement

The authors have declared no competing interest.

