## Supplementary Figures for "Neurofibromin 1 mediates sleep depth in *Drosophila*"

Supplemental Figure 1

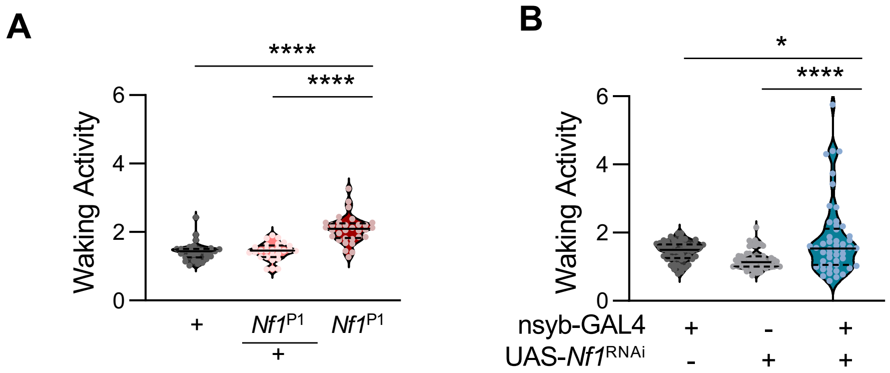

Supplemental Figure 2

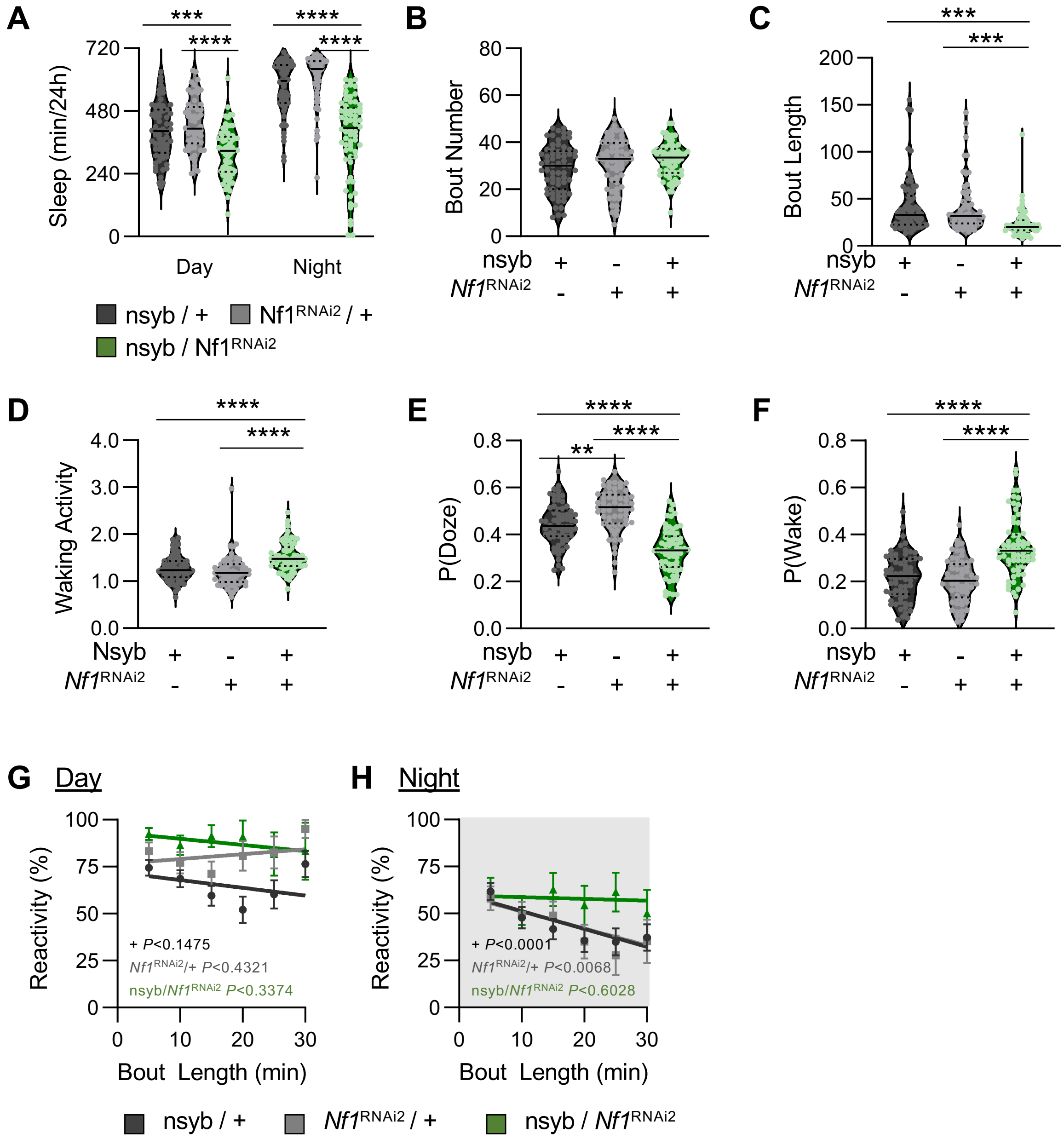

Supplemental Figure 3

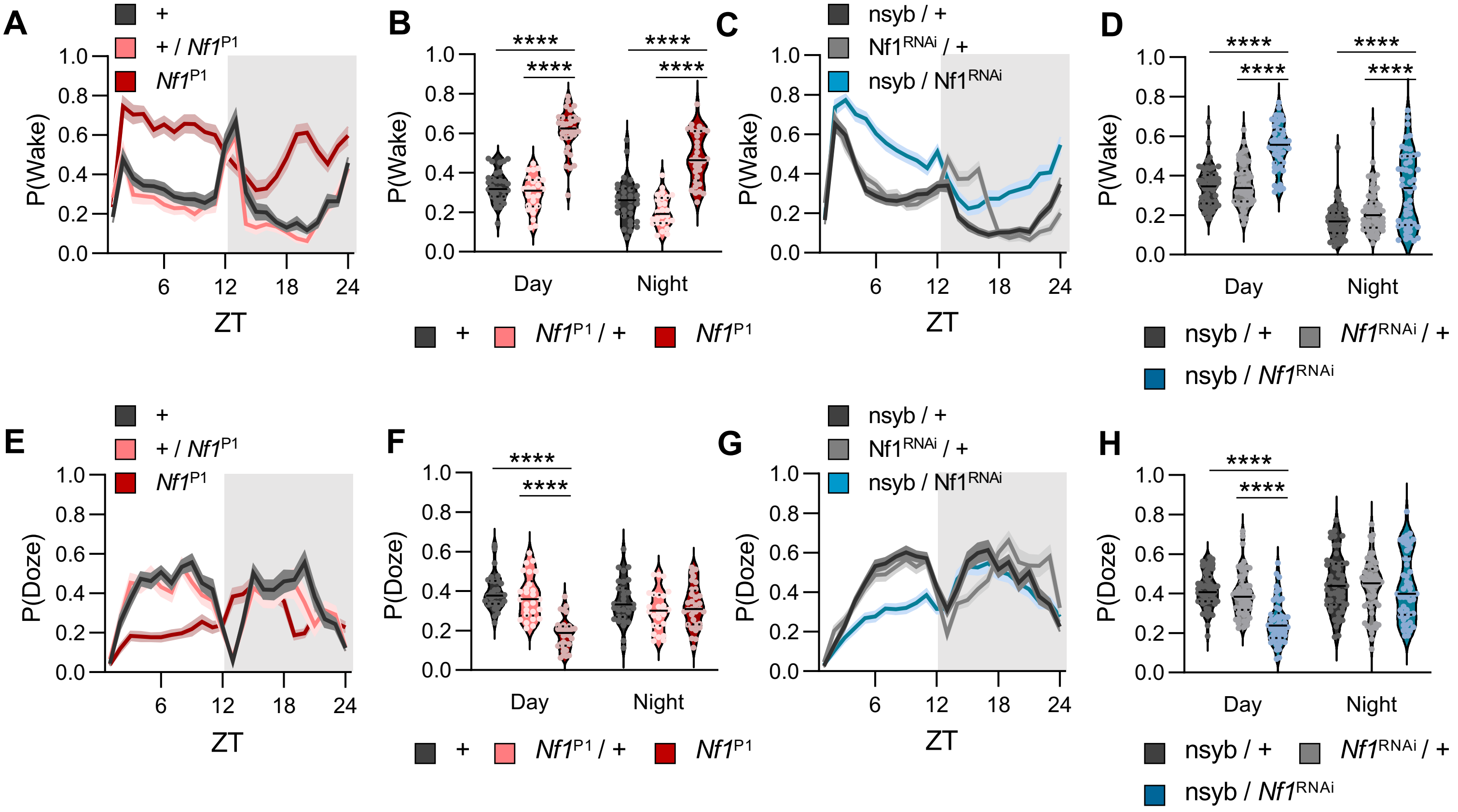

Supplemental Figure 4

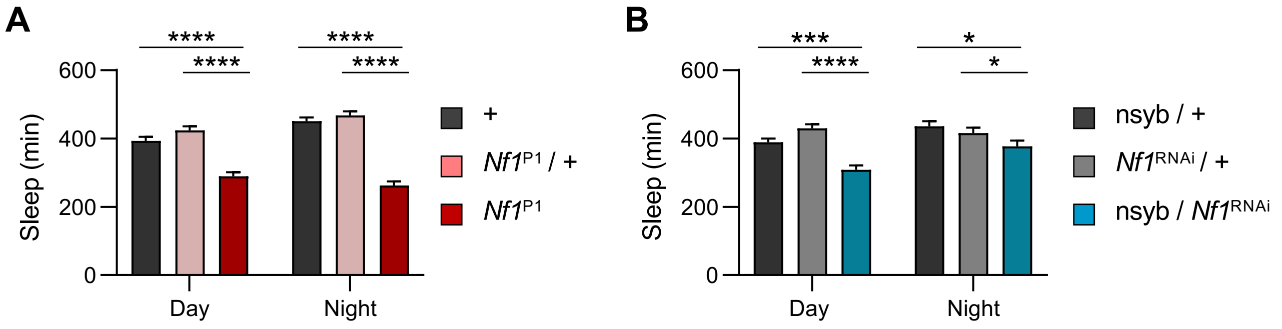

Supplemental Figure 5

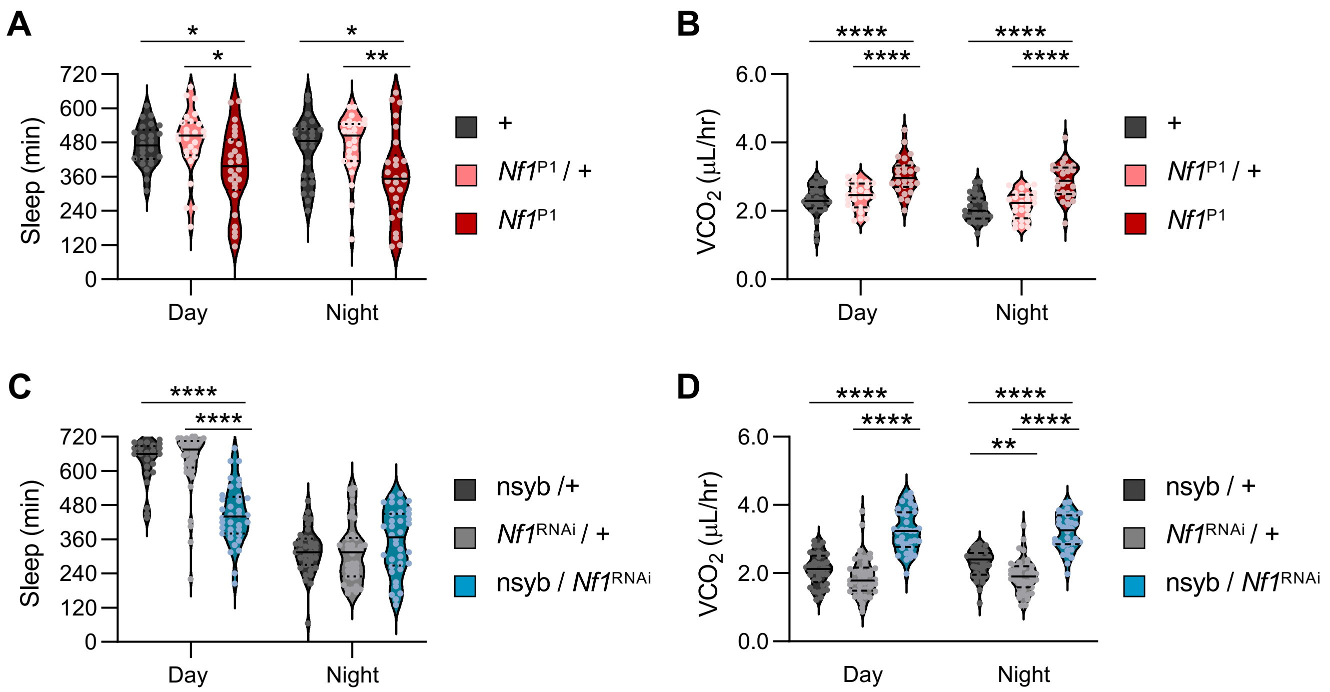

Supplemental Figure 6

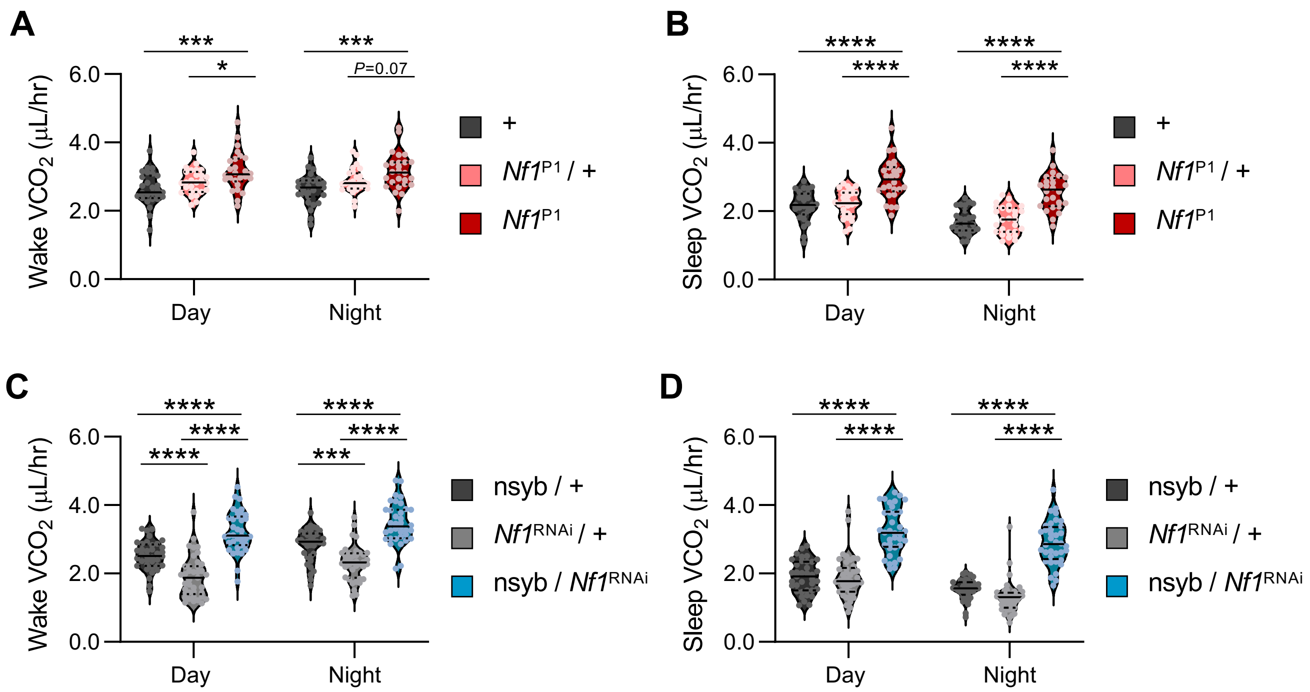

Supplemental Figure 7

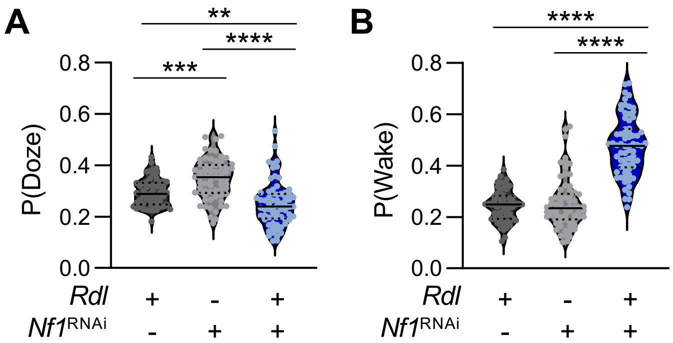

Supplemental Figure 8

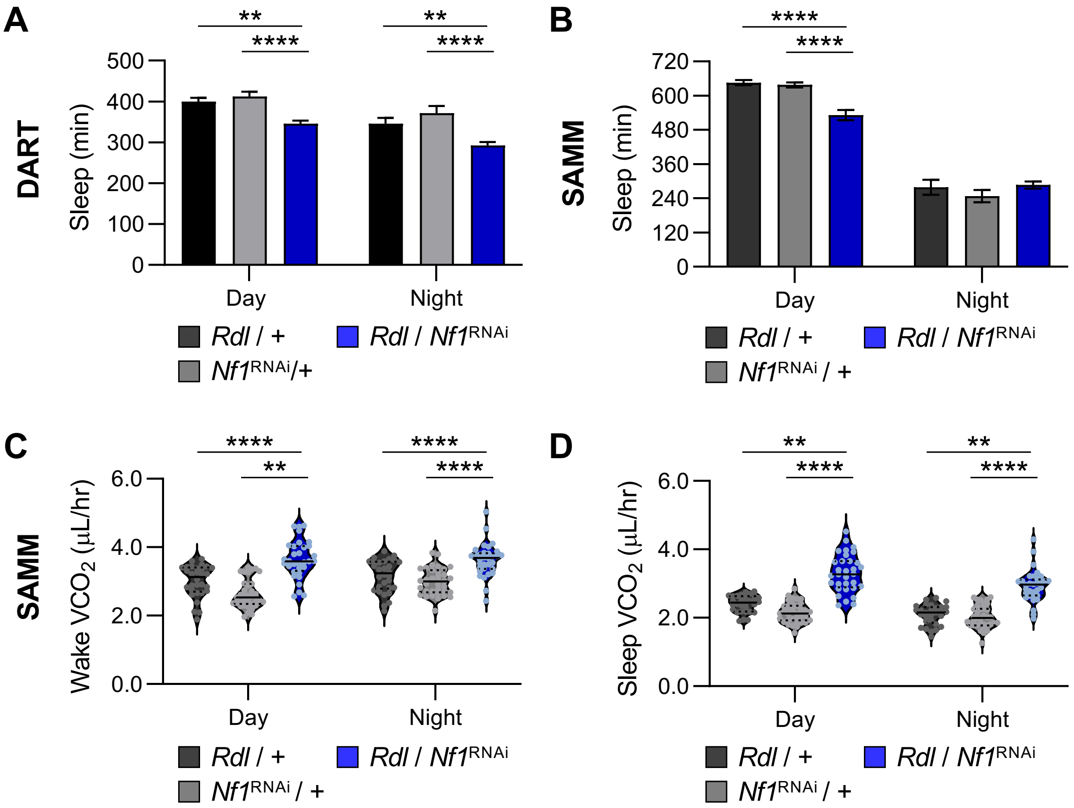

### Supplemental Figure 9

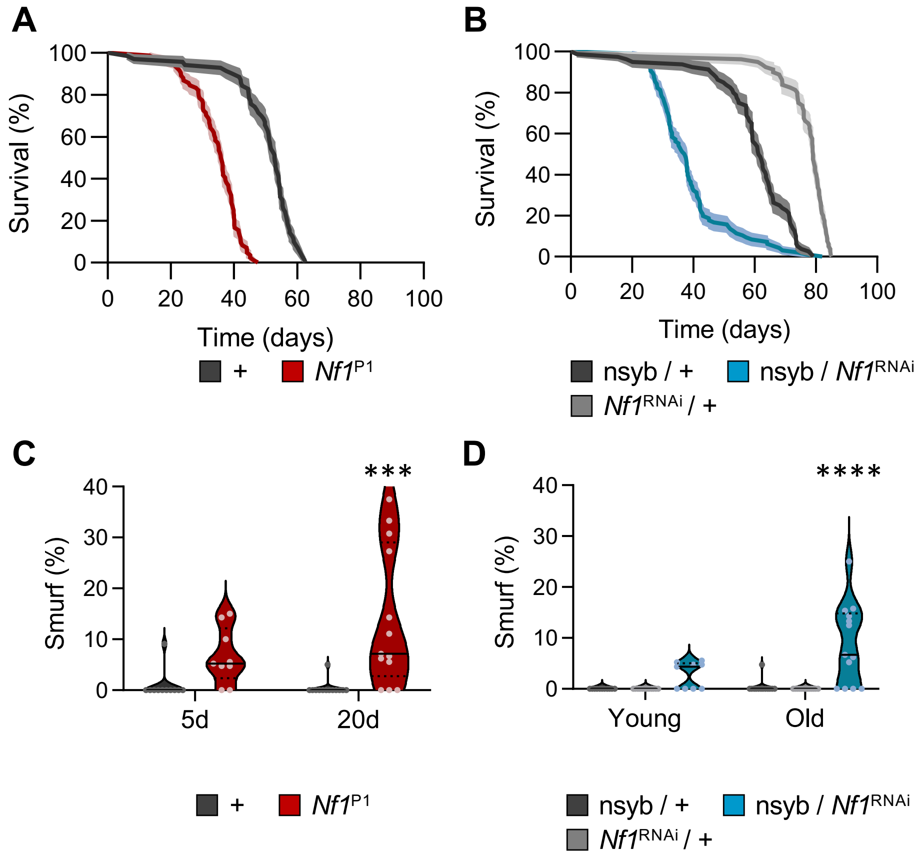

Supplemental Figure 10

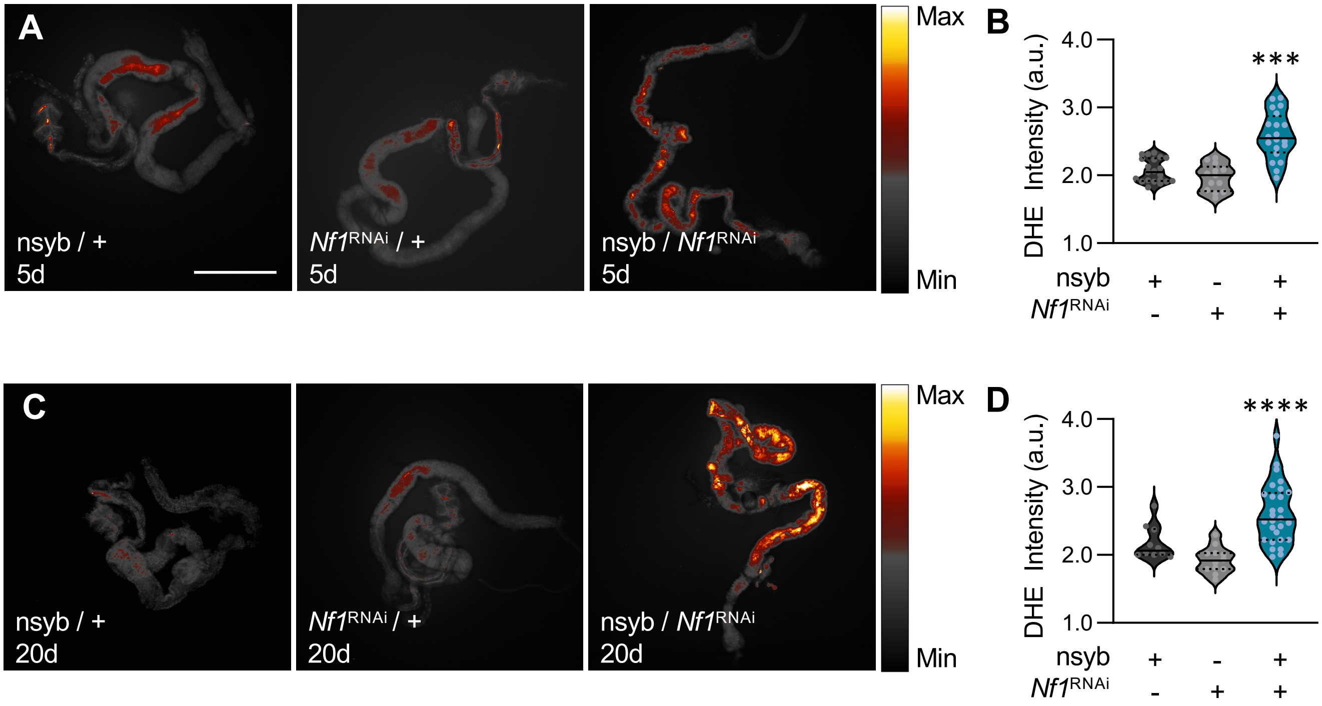
